# Molecular mechanism of cell-cell adhesion mediated by cadherin-23

**DOI:** 10.1101/208272

**Authors:** G. S. Singaraju, A. Kumar, J. S. Samuel, A. Sagar, J. P. Hazra, M. K. Sannigrahi, R. M. Yennamalli, Ashish, S. Rakshit

## Abstract

Adherin-junctions are traditionally described by the homophilic-interactions of classical cadherin-proteins at the extracellular region. However, the role of long-chain non-classical cadherins like cadherin-23(Cdh23) is not explored as yet even though it is implicated in tissue-morphogenesis, cancer, and force-sensing in neuronal tissues. Here, we identified a novel antiparallel-binding interface of Cdh23 homodimer in solution by combining biophysical and computational methods, *in-vitro* cell-binding, and mutational modifications. The dimer consists of two electrostatic-based interfaces extended up to two terminal domains, atypical to classical-cadherins known so far, and forms the strongest interactions in cadherin-family as measured using single-molecule force-spectroscopy. We further identified single point-mutation, E78K, that completely disrupts this binding. Interestingly, the mutation, S77L, found in skin cancers falls within the binding interface of the antiparallel-dimer. Overall, we provide the molecular architecture of Cdh23 at the cell-cell junctions which are likely to have far-reaching applications in the fields of mechanobiology and cancer.

## Introduction

Cadherin-23 (Cdh23) is abundantly expressed in nearly 90% of normal tissues (The Human Proteome Atlas) and 80% of cancer tissues (The Cancer Genome Atlas) and well-known for its role as a mechanosensor in neurosensory epithelial cells of the inner ear and retina. In the hair-cell stereocilia of the inner ear, Cdh23 interacts with protocadherin-15 (Pcdh15) to form tip-links that directly converts force from sound stimuli or head movement into electrical signals(Kazmierczak *et al*, 2007). During early development of the stereociliary organization and hair-bundle formation in hair-cells, Cdh23 is engaged in lateral links(Michel *et al*, 2005). In the retina, Cdh23 is localized at the synaptic region of the photoreceptive cells and mediates cell-cell adhesion(Lagziel *et al*, 2005). Truncation mutations and missense mutations in Cdh23 are responsible for Usher syndrome (USH1D), which causes hearing loss and progressive vision loss(Bork *et al*, 2001).

The Human Proteome Atlas (THPA) reported the high expression of Cdh23 in organs like brain, lymph node, kidney, gastrointestinal tract, testis and skin(Uhlén *et al*, 2015). Genome-wide association studies (GWASs) of kidney function have identified a strong association of Cdh23 with cross-sectional eGFR expression and its knockdown in zebrafish embryo resulted in severe edema, suggesting its important role in normal kidney functions(Gorski *et al*, 2015). The Cancer Genome Atlas (TCGA) data also showed increased expression of Cdh23 in most of the solid cancers including sarcoma, adrenocortical carcinoma, cervical cancer, head and neck squamous cell carcinoma, kidney renal clear cell carcinoma and breast cancer. Recent studies with the MCF-7 breast cancer cell-lines have confirmed the involvement of Cdh23 in homotypic cell-cell adhesion, similar to classical cadherins (Apostolopoulou & Ligon, 2012). Cdh23 was also observed at the heterotypic cell-cell junctions between MCF-7 and normal breast fibroblasts(Apostolopoulou & Ligon, 2012), a characteristic of early metastasis. In both these junctions, Cdh23 from apposing cell surfaces participate in homophilic interactions. *Ex-vivo* experiments with non-aggregating L929 cells expressing recombinant full-length Cdh23 were also shown to form aggregates exclusively via Cdh23-mediated homophilic interactions(Siemens *et al*, 2004). When admixed with another batch of L929 cells expressing recombinant full-length E-cadherin, the cells aggregated into two separate groups expressing either of the two proteins and did not form mixed aggregates(Siemens *et al*, 2004). These observations indicate that Cdh23 forms homophilic interactions, and heterophilic interactions with other non-classical cadherins (e.g., Pcdh15) but not with classical cadherins like E-cadherin. This sorting could be due to different binding mechanisms of the two classes of cadherins.

Cdh23 comprises a cytosolic domain, a single-pass transmembrane region followed by 27 extracellular (EC) domains, unlike just 5 EC domains in classical cadherins(Bork *et al*, 2001,Siemens *et al*, 2004,Söllner *et al*, 2004). However, similar to classical cadherins, Cdh23 causes cells to adhere using two types of homophilic interactions, *trans* and *cis*(Lagziel *et al*, 2005,Uhlén *et al*, 2015). Electron tomography revealed a unique pattern in the *cis*-homodimer of Cdh23: a pair of Cdh23 molecules aligned in the same orientation and intertwined to form a helical complex through interactions between all EC-domains except the two terminal ones, EC1 and EC2 (EC1+2) which remained exposed outward to facilitate *trans*-interactions(Kazmierczak *et al*, 2007). The engagement of these two terminal domains in a heterophilic trans-interaction was already reported with Pcdh15(Sotomayor *et al*, 2012). While the crystal structure of the Cdh23-EC1+2 monomer is known, no reports have revealed the molecular details of the homophilic *trans*-dimerization of Cdh23.

Classical cadherins undergo homophilic *trans*-interactions using the outermost terminal domain, EC1(Brasch *et al*, 2012, Sivasankar *et al*, 2009). They first form an X-dimer, a kinetically driven interaction via the overlap of the linker region between two terminal domains(Rakshit *et al*, 2012,Manibog *et al*, 2016). Eventually, the X-dimer is converted to a more thermodynamically stable strand-swap dimer (S-dimer) via an intermediate. The S-dimer is formed by swapping the N-terminal β-strand, containing a conserved tryptophan at position 2 (W2) or 4 (W4), to a hydrophobic core of the EC1 domain of the binding partner(Ciatto *et al*, 2010,Harrison *et al*, 2010). Among non-classical cadherins, desmosomal cadherins form an S-dimer(Garrod *et al*, 2005) whereas T-cadherin and R-cadherin form an X-dimer(Ciatto *et al*, 2010). Cdh23, however, lacks the sequence-determinants for either S- or X-dimerization. Moreover, the EC1 domain of Cdh23 has several unique features: a 5 residue long 3_10_-helix just prior to the A* β-strand, a α-helical loop connecting two β-strands and most strikingly, an additional Ca^2+^-binding site toward the N-terminus(Sotomayor *et al*, 2010). Together, these features indicate a unique interface for Cdh23-mediated homophilic *trans*-interactions.

We observed two distinct binding interfaces in the Cdh23-EC1+2 homophilic trans-dimer that exists in an extended handshake conformation. This conformation is architecturally similar to the heterophilic *trans*-dimer between Cdh23 and Pcdh15, found at the tip-links(Sotomayor *et al*, 2012). However, the residues involved in the interactions differ. Interface one in the Cdh23-EC1+2 *trans*-homodimer involves residues of EC1 alone. Interestingly, EC1 alone did not form any heterodimer with Pcdh15(Sotomayor *et al*, 2012). The second interface is formed by the interactions between residues from both EC1 and EC2. The involvement of EC2 in homophilic *trans*-interactions is rare in cadherin family. Both interfaces thus jointly form a stable complex with high-affinity, as measured using single-molecule force spectroscopy (SMFS). To the best of our knowledge, this is the strongest interaction ever measured for homophilic adhesion between any cadherin(Panorchan, 2006,Bajpai *et al*, 2008).

## Results

### Cdh23-EC1+2 is involved in Cell-cell adhesion

Cdh23 exists in 9 different variants (NG_008835.1). Variant 1 is a full-length cadherin, containing all 27 EC domains, a transmembrane domain, and cytosolic component. Variants 2,3,4,5 vary in length yet have proximal EC domains intact, whereas variants 6 and 7 contain only last 7 EC domains. Variants present in MCF-7 cell-lines were identified from the mRNA expression using qRT PCR (Supplementary Fig. 1a) and protein expression using Western blot (WB) (Supplementary Fig. 1b). Both studies suggested the presence of full-length Cdh23 (variant 1) along with variants 2,3,4,5 and all contain EC1+2 domains. This was expected as EC1+2 domains only can form homophilic trans-interactions and it corroborates with the previous studies where the use of functional antibodies against EC1 domain of Cdh23, weakened the adherent junctions in MCF-7. We could not detect the presence of variant 6 and 7 using qRT PCR as these variants have complete sequence match with variant 1, however, we observed the expression of these variants in WB (Supplementary Fig. 1b).

### Cdh23-EC1+2 forms a homodimer in solution

In order to understand the molecular details of Cdh23-mediated trans-homodimer, we expressed Cdh23-EC1+2 in *E. coli BL21RIPL* following a reported protocol (Sotomayor *et al*, 2010). The expressed protein was purified in two steps by Ni^2+^-NTA-based affinity followed by size-exclusion chromatography (SEC) (Materials and Methods). To assess the formation of higher-order structures of Cdh23-EC1+2 in solution, we concentrated the protein to 130-150 μM and performed SEC. Two peaks at the elution volumes of 15.6 mL (P1) and 16.9 mL (P2) were observed in the chromatogram (Column volume: 24 mL) (Fig. 1a). We subsequently ran SDS-PAGE of all the eluted fractions and observed a single band corresponding to 26 kDa (Fig. 1a, inset, Supplementary Fig. 2), suggesting that P2 and P1 corresponded to the Cdh23-EC1+2 monomer and a higher-order association, respectively. To determine the molecular weights of the elutions, we developed a calibration curve by running SEC with standard proteins of varying molecular weights under the same conditions (Supplementary Fig. 3). The first and second peaks corresponded to the molecular weights of 25 and 50 kDa, respectively; suggesting the higher-order association as a dimer.

**Figure 1.**
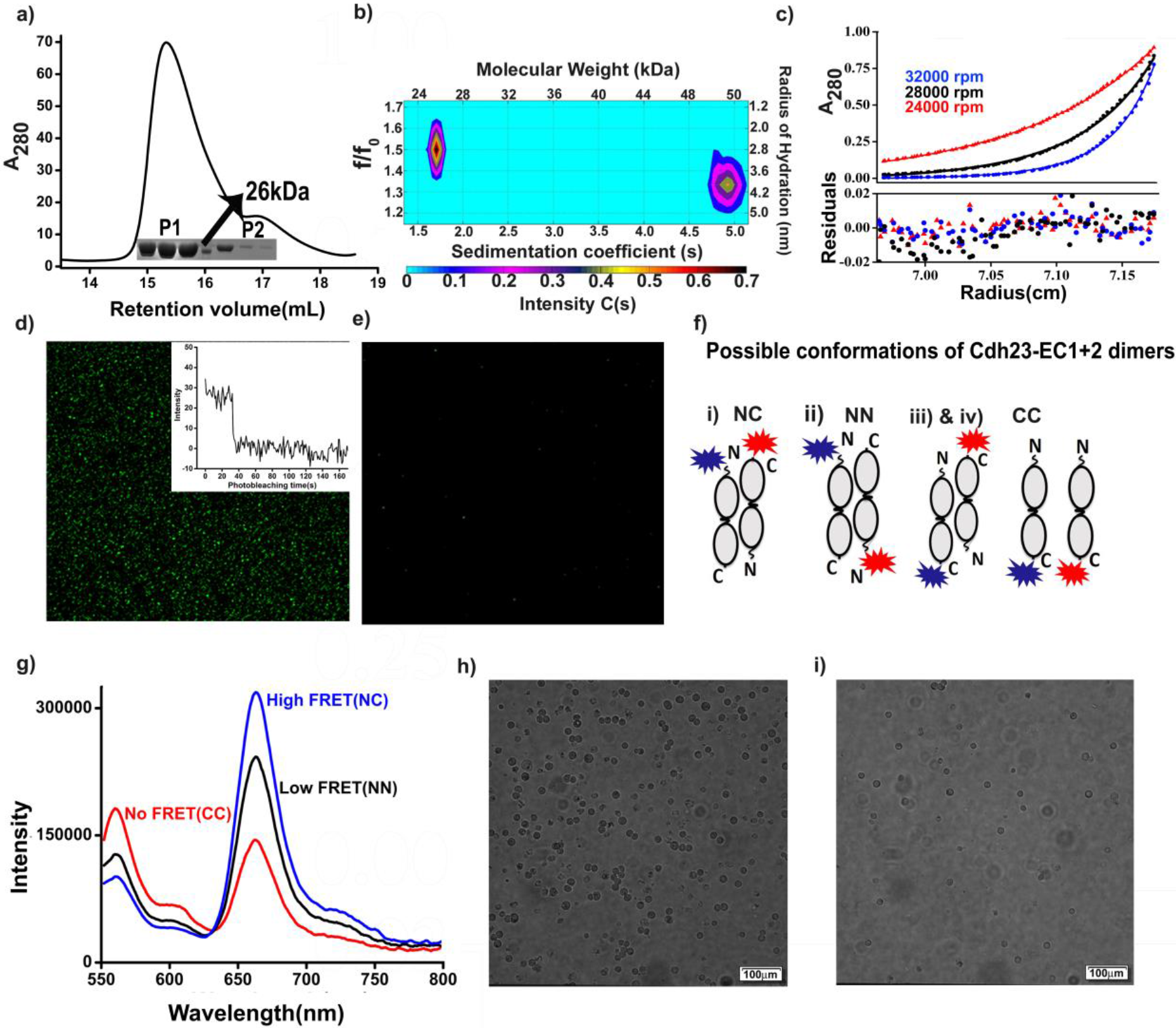
Homodimerization of cadherin-23 in the *trans*-conformation. **a)** Analytical SEC of 130 μM Cdh23-EC1+2 showed two peaks with respect to the elution volume. The first peak, eluted at 15.6 mL, corresponded to a dimer, and the second peak, eluted at 16.9 mL, corresponded to a monomer. The inset shows the SDS-PAGE of the equal fractions of elution. The observed single band at position ~26 kDa corresponded to the theoretical estimate for Cdh23-EC1+2. **b)** The SV profile of Cdh23-EC1+2 is presented with a color map to show the correlations between the sedimentation coefficients (s), frictional coefficients (f/f_0_), radii of hydration and molecular weights of the populations. The two populations in the profile confirmed the presence of dimers and monomers in solution when running at a concentration of 36 μM at 20°C. **c)** SE experiments of Cdh23-EC1+2 were run for 40 μM protein at three rotor speeds, 24,000 (red dots), 28,000 (black dots) and 32,000 rpm (blue dots). The red lines represent the global fitting of the data to the monomer-dimer equilibrium model using SEDPHAT. The residuals of each fit are displayed below. **d)** Fluorescence images showing the pull-down signal for Cdh23-EC1+2 WT at 2mM Ca^2+^. The green spots represent the proteins being pulled from solution by the Cdh23-EC1+2 proteins attached to coverslips, via homodimerization. The inset shows the representative single-step photobleaching of each fluorophore attached to proteins being pulled. **(e)** The signal was lost after washing the coverslip with EGTA. **f)** Scheme showing all the possible conformations of dimers and naming the constructs as NC, NN, and CC based on whether the donor and acceptor dyes were attached to the N-terminal and C-terminal, respectively. **g)** Steady-state fluorescence showed stronger emission of cy5 with the excitation of cy3 (λ_*ex*_ = 545 nm) for NC than for NN, suggesting higher FRET for NC than for NN. Almost negligible FRET was observed for CC with same excitation (λ_*ex*_ = 545 nm). For all experiments, the concentration of each dye-modified protein was maintained at 55 μM. **h)** Live Breast cancer cell-lines, MCF-7, were found adhered to coverslips coated with Cdh23-EC1+2. **i)** Washing with EGTA to chelate Ca^2+^ - ions, detached cells from the surface.

The higher-order association corresponds to a dimer, as determined by dynamic light scattering (DLS), as well. The obtained hydrodynamic radius (*R*_*H*_) in the presence of Ca^2+^ ions was 3.9 nm (Supplementary Fig. 4), nearly double that of the monomer (2.2 nm) as estimated from the crystal structure (Eq. 1 in SI). Upon removal of Ca^2+^ ions either by chelation with ethylene glycol-bis (β-aminoethyl ether)-N,N,N’,N’-tetraacetate (EGTA) or by washing through a chelex column, the *R*_*H*_ of the protein was reduced to 2.7 nm, corresponding to the monomer (Supplementary Fig. 4). This suggested that similar to classical cadherins(Siemens *et al*, 2004), Cdh23-EC1+2 undergoes Ca^2+^-dependent dimerization. Similar Ca^2+^-dependent adhesion was observed in fluorescent-based single-molecule pull-down assays on a glass-coverslip (Fig. 1d, e).

We performed sedimentation velocity (SV) experiments using analytical ultracentrifugation (AUC) at two protein concentrations, 10 μM and 36 μM, with a rotor speed of 42,000 rpm at 20°C. The lower concentration run showed one dominating peak with a sedimentation coefficient of 1.96 s and a frictional coefficient ratio (f/f_0_) of 1.51, indicating a monomer with a molecular weight of 26 kDa and *R*_*H*_ of 2.8 ± 0.1 nm (Supplementary Figs. 5 and 6). The SV run at the higher concentration produced two distributions of C(s). The peak that appeared at 2.1 s corresponding to the monomer, while the peak at 4.77 s and lower f/f_0_) (1.34) corresponded to the dimer (Fig. 1b). Further, we confirmed the existence of the dimer from the estimation of the molecular weight (52 kDa) and R_*H*_ (4.1±0.3 nm). The R_*H*_ obtained from SV corroborated that obtained from DLS. We then performed sedimentation equilibrium (SE) using AUC with 40 μM protein (and 60 μM) at three different rotational speeds (24,000 rpm, 28,000 rpm and 32,000 rpm) and estimated the *K*_*D*_ of monomer-dimer equilibrium as 18 ± 4 μM from the global fitting of the equilibrium curves using free software SEDPHAT(Fig. 1c)(Schuck, 2003).

### The homodimer of Cdh23-EC1+2 is a *trans*-dimer interacting via N-termini

Next, we designed ensemble-FRET to understand the orientation of the constituent monomers in the dimer. For FRET experiments, proteins were labeled with the FRET-pair dyes cyanine-3 (cy3) and cyanine-5 (cy5) (Fig. S6), at the C-terminus and N-terminus, respectively (Supplementary Fig. 7 and Materials and Methods)(Guimaraes *et al*, 2013). To avoid artifacts due to dyes, we repeated the experiments with another set of dyes, IAEDANS as donor and fluorescein maleimide as acceptor (Supplementary Fig. 8). The dye to protein labeling ratio was 1:1. Three FRET experiments were performed: 1) C-terminal cy3-labeled protein was added to the C-terminal cy5-labeled protein (referred to as the C-C construct); 2) In N-N construct, N-terminal cy3-labeled protein was added to N-terminal cy5-labeled protein; and 3) C-terminal cy3-labeled protein was added to N-terminal cy5-labeled protein (C-N construct). The possible conformations that could form from these protein mixtures are shown schematically in Fig. 1e. The C-C construct showed no FRET (Fig. 1f). In contrast, both the N-N and C-N constructs showed positive FRET with two different FRET efficiencies of 0.64 for the C-N construct and 0.15 for the N-N construct (Fig. 1f and Supplementary Table 3). Similar trends were observed in the lifetime-decays of the donors (Supplementary Fig 9). Since FRET efficiency is inversely proportional to the distance between FRET pairs, we conclude that the C- and N-termini of the opposing partners in the homodimer are closer than the N-N termini, forming an extended overlap.

Further, to check the physiological relevance of this dimer-conformation and the ability of Cdh23 EC1+2 forming cell-cell adhesion, we performed *in-vitro* binding of MCF-7 live-cells on Cdh23 EC1+2 modified surfaces. This assay was based on the facts that MCF-7 cells endogenously express predominantly two types of cell-cell adhesion proteins, Cdh23 and E-cadherin, and these proteins do not interact with each other(Apostolopoulou & Ligon, 2012). We, therefore, attached the C-terminus of Cdh23 EC1+2 covalently onto the glass-coverslips using the sortagging protocol(Srinivasan *et al*, 2017) and allowed ~10^6^ MCF-7 live-cells to interact to the protein-coated coverslips for 45 minutes. After vigorous washing with Ca^2+^-buffer, we observed a large number of cells remained adherent to the surface (Fig. 1g). However, cells became non-adherent in presence of EGTA (Fig. 1i). These findings indicate that the EC1+2 domains are enough to mediate Ca^2+^ - dependent cell-cell adhesion by *trans*-interactions via the N-termini.

### SAXS envelope suggests a compact extended-handshake conformation for the dimer

To reveal the shape of the Cdh23-EC1+2 *trans*-homodimer in solution, we performed SAXS measurements with proteins at 10mg/mL (Fig. 2,a and b), for a q-range of 0.1 to 4 nm^−1^ using SAXSpace instrument (Anton Paar GmbH, Austria) (Table S4). The monodispersity of the sample was confirmed from the linearity of the Guinier plot (Fig. 2a, inset), and the proper folding of the proteins from a strong agreement between the normalized Kratky plots (I(q)*(q*R_g_)^2^/I(0) vs q*R_g_) of the experimental data and the theoretical SAXS profiles of the crystal structures/models. It is pertinent to mention here that while the peaks of the normalized Kratky plots deviate from the value of 1.73, this can be attributed to the rod-like nature of the protein, as the theoretical SAXS profile of the docked model also showed the same profile. (Fig. 2b, inset)(Konarev *et al*, 2003). From the indirect Fourier transformation of the SAXS profile using GNOM (Svergun, 1992), we generated the pair-distribution function (P(r)), which provided the frequency of the interatomic vectors inside the predominant shape of the proteins in real space(r). P(r) curves estimated for dimer showed maximum linear dimension (D_max_) and R_g_ values of 11.6 and 3.2 ± 0.2 nm, respectively (Fig. 2b). The peak position and shoulders profile of the computed P(r) supported the multi-domain shape of Cdh23 connected by a non-flexible linker.

**Figure 2.**
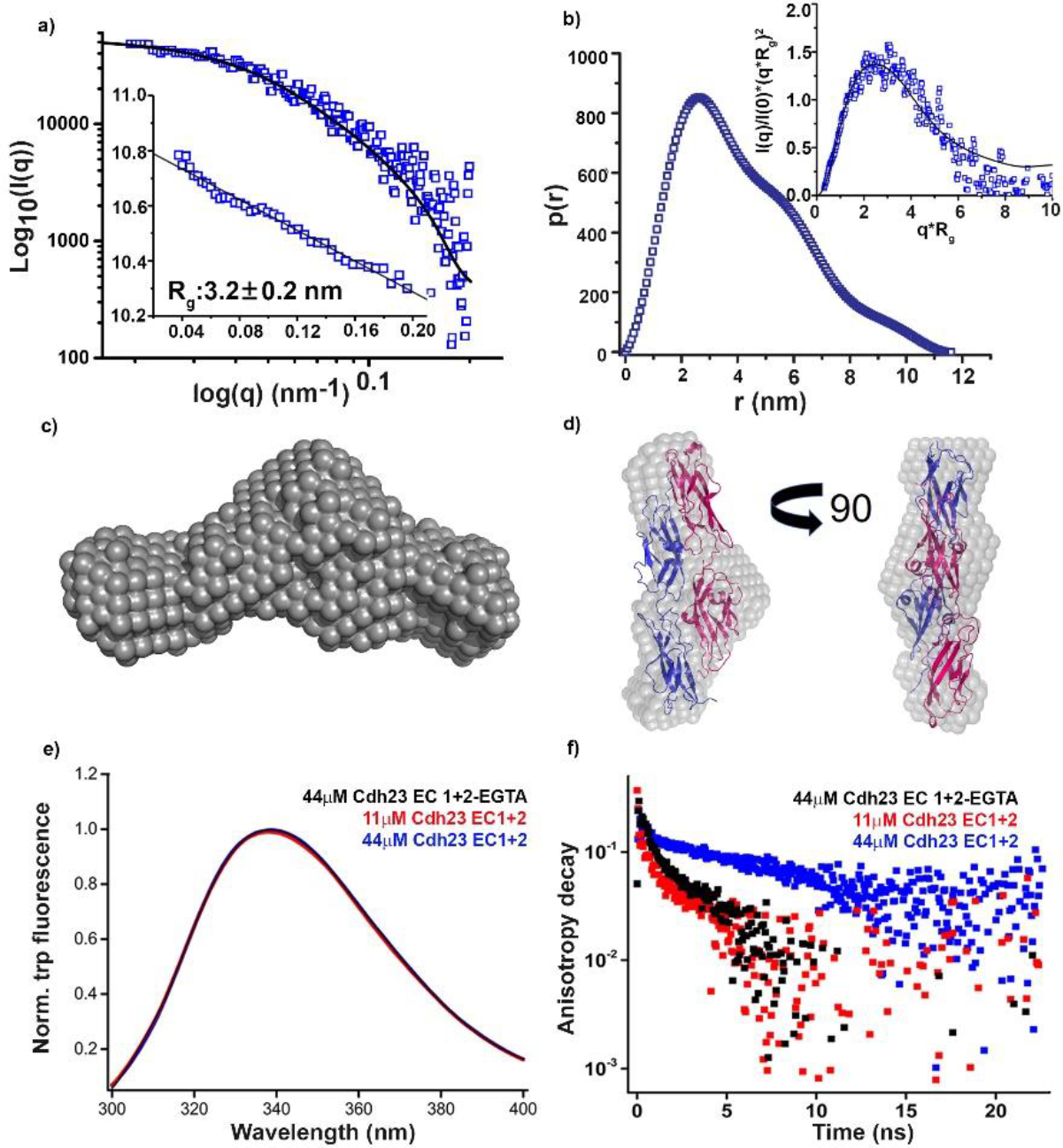
Cadherin-23 dimer forms an extended handshake conformation. **a)** SAXS profile of the Cdh23-EC1+2 dimer was plotted as the scattering intensity I(q) vs. the scattering vector (q) on a log-log scale. The solid black line represents the theoretical SAXS profile computed from the proposed model of the dimer. The linear nature of the Guinier plot (ln(I(q)) vs. q^2^) for the low q region is marked (inset, bottom, left). **b)** The p(r) distribution for the dimer (blue) showing the maximum linear dimensions (*D*_*max*_) to be 11.6 nm. The normalized Kratky plot (I(q)/I(0)*(q*R_g_)^2^ of the experimental SAXS profile superimposed with that of the theoretical SAXS profile of the proposed dimer is shown in the inset. **c)** The average *ab-initio* model obtained from DAMMIN is shown for the dimer. **d)** The statistically verified and rank-1 patch-dock structure (PDB: 2WHV) of the dimer (blue) is fitted to the SAXS envelope (gray) for the dimer and shown with two orthogonal perspectives. **e)** Steady-state fluorescence was measured for Cdh23-EC1+2 (λ_ex_ = 295 nm; λ_em_ = 342 nm) to identify the change in the W66 environment in different protein states: monomeric (blue & black) and dimeric (red). **f)** Time-resolved fluorescence anisotropy decays of *W66* in Cdh23-EC1+2 were recorded for the monomer (red & black) and dimer (red) to track its role in dimerization.

To visualize the three-dimensional shape of the dimer, ten independent dummy residue models were generated using DAMMIF(Franke & Svergun, 2009), averaged using DAMAVER, refined again using DAMMIN(Svergun, 1999), and compared with each other by calculating the normalized spatial discrepancy (NSD). NSD is a measure of the similarity in the shapes of the models with a value of < 1 indicating that the models are fairly similar and values above 1 indicating that the models are systematically different from each other. The mean NSD between the 10 models was 0.602 with a standard deviation of 0.027 indicating that all the models were very similar to each other and the modeling protocol was stable and reproducible. The resulting envelope had dimensions of 14(L)×5.76(W)×4.1(H) nm (Fig. 2c).

### Cdh23-EC1+2 homodimerization is not mediated by tryptophan

To identify a dimer complex that could fit the SAXS envelope, we first considered the W-conformation proposed previously(Sotomayor *et al*, 2010). It was predicted that Cdh23-EC1+2 might form a trans-homodimer through π-stacking of the indole ring of the sole tryptophan at 66 (W66) (Supplementary Fig. 12). The driving force behind such W-mediated interactions is the switch of the W-environment from hydrophilic to more hydrophobic. Since W is known to feature a solvatochromic shift in fluorescence, we designed experiments to probe the W-emission of the monomer and dimer and decipher its role in dimerization. Accordingly, we monitored the steady-state emission of W66 (λ_ex_ = 295 nm) as the protein concentration increased from a monomer concentration to beyond the *K*_*D*_ (~18 μM) of the dimer (Fig. 2e). We did not observe any shift in W-emission, excluding its possible involvement in dimerization. Further, we examined the mobility of W66 in the monomer and dimer using time-resolved fluorescence anisotropy. W in a protein can have two rotational components, a fast rotation around its own axis and a relatively slower rotation along with the protein. Dimerization via the π-stacking of the W residues is expected to constrain the free local rotation around its axis and therefore delay the anisotropy decay of the fast component. We probed the anisotropy decay of W66 at concentrations ranging from monomer to dimer (Fig. 2f) and observed no change in the decay rate of the faster components. This indicated a lack of constraints on the local mobility of W66 and thus confirming that W66 did not play a role in dimerization. However, we observed a significant change in the decay for the global rotation of W66 upon dimerization, as expected (Fig. 2f and Supplementary Table 6).

We then searched for new structures by docking Cdh23-EC1+2 (PDB ID: 2WHV) using PatchDock. Of the first ten models based on dock-scoring, seven models showed docking through the N-terminus of the protein and the others through the C-terminus. We considered these seven models and superimposed them with the SAXS envelope using SUPCOMB, which showed comparable NSD values varying from 0.7–0.8 for all the structures. However, rank 1 fit best from the z-tests based on R_g_ and FRET-based end-to-end distances (p = 0.20) (Fig. 2d and Supplementary Table 7).

We next performed molecular dynamics (MD) simulations of the rank 1 structure using GROMACS and delineated residue-wise interactions (Materials and Methods, Supplementary Table 8). Stability of the dimer was achieved within 2 ns of the simulations to an average root mean square deviation (RMSD) of 0.34 nm (Fig. 3a and b). To identify the residues that are involved in dimerization, we compared the root mean square fluctuations (RMSF) of all residues between the monomer and dimer anticipating that residues buried within interacting surfaces may fluctuate less (Fig. 3c). The residues other than glycines that showed significant fluctuation differences were highlighted (Fig. 3c, inset). The time-trace analysis of the MD results also indicated these residues responsible for the interactions. We divided these residues into two main interacting interfaces based on their positions in the EC domains (Fig. 3, a-g). Interface one is dominated by electrostatic interactions that are mediated by the antiparallel overlap of strand F of the EC1 repeats. The residues involved in the interactions are K76, S77, E78, N97 and Q99, which are conserved in human, mouse, rat, and zebrafish (Supplementary Fig. 13). Interestingly, S77 was found mutated to L in patients suffering from skin cutaneous cancer. Some of these residues are also instrumental in heterodimerization with Pcdh15 at tip-links. The other interface, which is amphiphilic in nature, is formed by anchoring of the elongated N-terminal strand of EC1 of one monomer to the β-strand (G-strand, Supplementary Fig. 14) of EC2 of the other. None of these residues are occluded by glycosylation, indicating that the interfaces are physiologically relevant (Supplementary Fig. 15). This was evident from the *in-vitro* cell-binding assays as well, which showed the arrest of MCF-7 cell-lines on surfaces coated with bacterially expressed Cdh23-EC1+2 with no translational modifications. Further, we introduced a single point mutation at E78 to a positively charged residue (E78K). Proper folding of the mutants was verified using SEC and circular dichroism (Supplementary Figs. 10 and 11). E78K showed complete disruption of the dimer. The *R*_*H*_ of E78K obtained from the SV run was 2.9 ± 0.1 nm (Fig. 4a and Supplementary Fig. 16) which is equal to the R_g_ (Fig. 4b) obtained from SAXS, corresponding to a monomer. The indirect Fourier transform gave a Dmax of 10 nm (Fig. 4c). These values correlated well with the R_g_ and *D*_*max*_ of 2.9 and 10.0 nm, respectively obtained from the X-ray crystal structure of protein (PDB ID: 2WHV). Further, we were able to fit the SAXS envelopes (Fig. 4d) obtained for E78K to the crystal structure of the monomer and the docked structure by aligning their inertial axis using SUPCOMB (Fig. 4e). Interestingly in the live-cell binding assays, MCF-7 cells were able to bind to surfaces modified with E78K. This may be due to weak interactions between wildtype (WT) proteins from the cell-surfaces to the mutants on coverslips.

**Figure 3.**
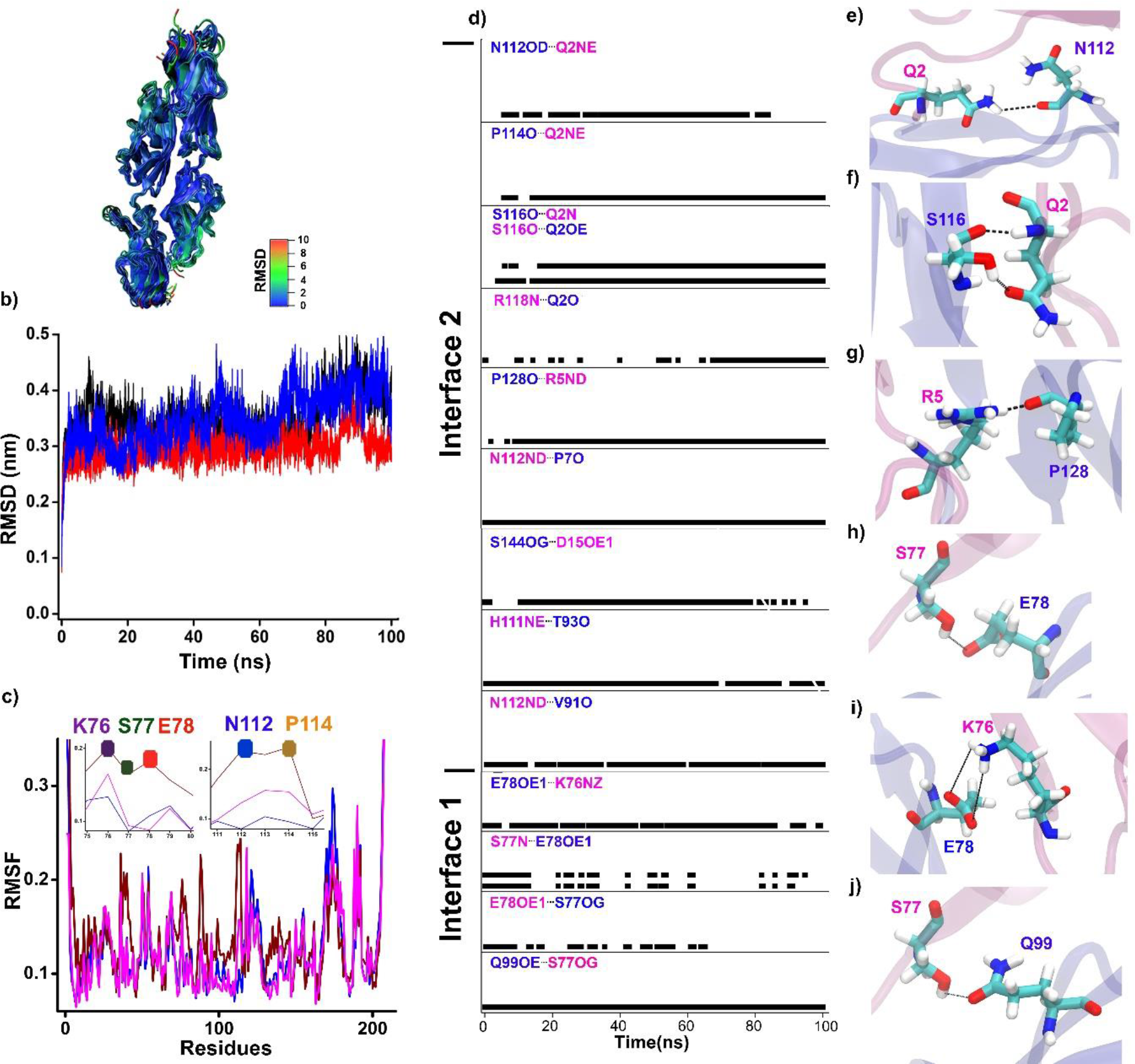
Survey of the H-bonds that are crucial for dimerization. **a)** Superimposed snapshots of the Cdh23-EC1+2 dimer at 10 ns intervals of the entire trajectory of the MD run are shown as a visual guide to the RMSD plot. **b)** RMSD the rank-1 docked Cdh23-EC1+2 dimer is shown for 3 independent MD runs of 100 ns. **c)** The RMSF is shown to compare the average fluctuations of all residues in chain A (magenta) and chain B (blue) of the dimer with the monomer (wine red). The zoomed plots in the inset highlight the RMSF of the interacting residues (left: K76 (purple), S77 (green), E78 (red) from interface 1 and right: N112 (blue) & P114 (yellow) from interface. **d)** Survival probabilities of the H-bonds over time are shown between different residues. From bottom to top, the interaction residues are divided into interface 1 and interface 2. Interface 1 contains residues within the EC1 domain alone, whereas residues from both EC1 and EC2 are involved in interface 2. The residues in pink are from chain A and the blue residues are from chain B of the dimer. **e-j)** The relative position of the residues involved in H-bonding are shown in enlarged diagrams between two chains, Chain A(magenta) and Chain B(blue): **e)** H-bonds between N112-Q2; **f)** Q2-S116; **g)** R5-P128; **h)** E78-S77; **i)** K76-E78 and **j)** Q99-S77. The represented H-bonds are highlighted with black dots.

**Figure 4.**
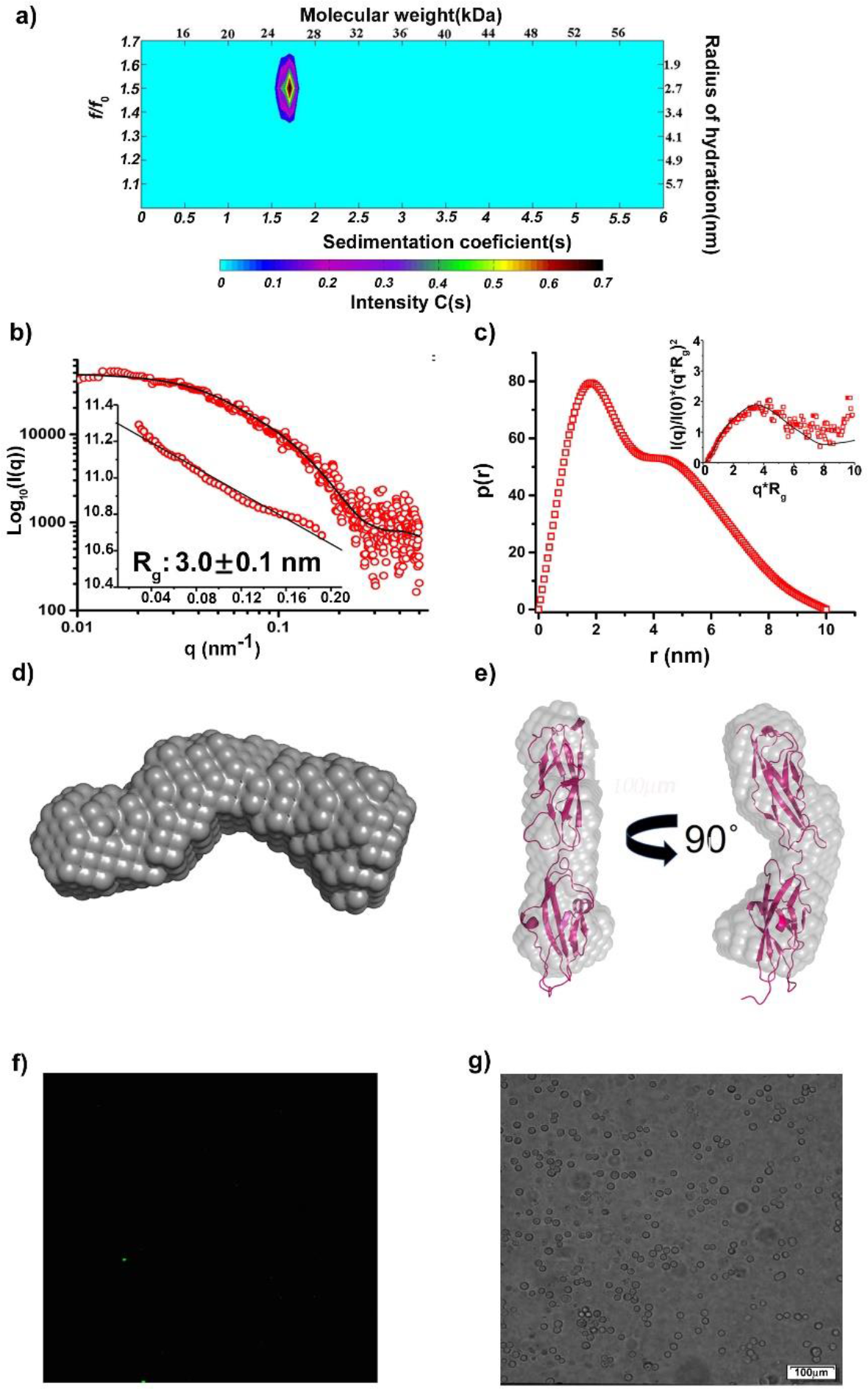
An E78K mutant of Cdh23-EC1+2 does not interact to form *trans*-homodimer. **a)** The SV profile of Cdh23-EC1+2 E78K is presented in color map to show the correlations between the sedimentation coefficients (s), frictional coefficients (f/f_0_), *R*_*H*_ and molecular weight. **b)** SAXS profile of the Cdh23-EC1+2 mutant E78K was plotted as intensity I(q) vs. scattering vector (q) on a log-log scale. The black solid line represents the theoretical SAXS profile computed from the crystal structure. The Guinier plot (ln(I(q)) vs. q^2^, q in nm^−1^) for the low q region is shown in the inset. **c)** The p(r) distribution for the E78K shows the maximum linear dimensions (*D*_*max*_) to be 10.0 nm. The normalized Kratky plot (I(q)/I(0)*(q*R_g_)^2^ of the experimental SAXS profile superimposed with that of the theoretical SAXS profile of the crystal structure is shown in the inset. **d)** The average *ab-initio* model obtained from DAMMIN is shown. **e)** For a visual guide, the averaged model is fit to the Cdh23-EC1+2 WT monomer (PDB ID:2WHV) using SUPCOMB. **f)** Fluorescence images do not show any signal for the proteins being pulled from solution by the Cdh23-EC1+2-E78K proteins attached to coverslips due to a loss in binding via homodimerization. **g)** Live Breast cancer cell-lines, MCF-7, were found adhered to coverslips coated with Cdh23-EC1+2-E78K.

Since interface one is mediated by EC1 alone, we checked whether the interface one interacts independently to form the homodimer by truncating EC2 from Cdh23-EC1+2. As expected, SV with Cdh23-EC1 (Supplementary Fig. 16) showed a population of both monomers and dimers indicating that EC1 alone can form a homodimer independently.

### The Cdh23-EC1+2 *trans*-homodimer is long-lived

To measure the strength of the interfaces in the dimer, we performed SMFS with Cdh23-EC1+2. We covalently attached the C-terminus of the protein to AFM cantilevers (Si_3_N_4_) and glass-coverslips using the sortagging protocol (Srinivasan *et al*, 2017) (Fig. 5a, Supplementary Fig. 17, Materials and Methods).

**Figure 5.**
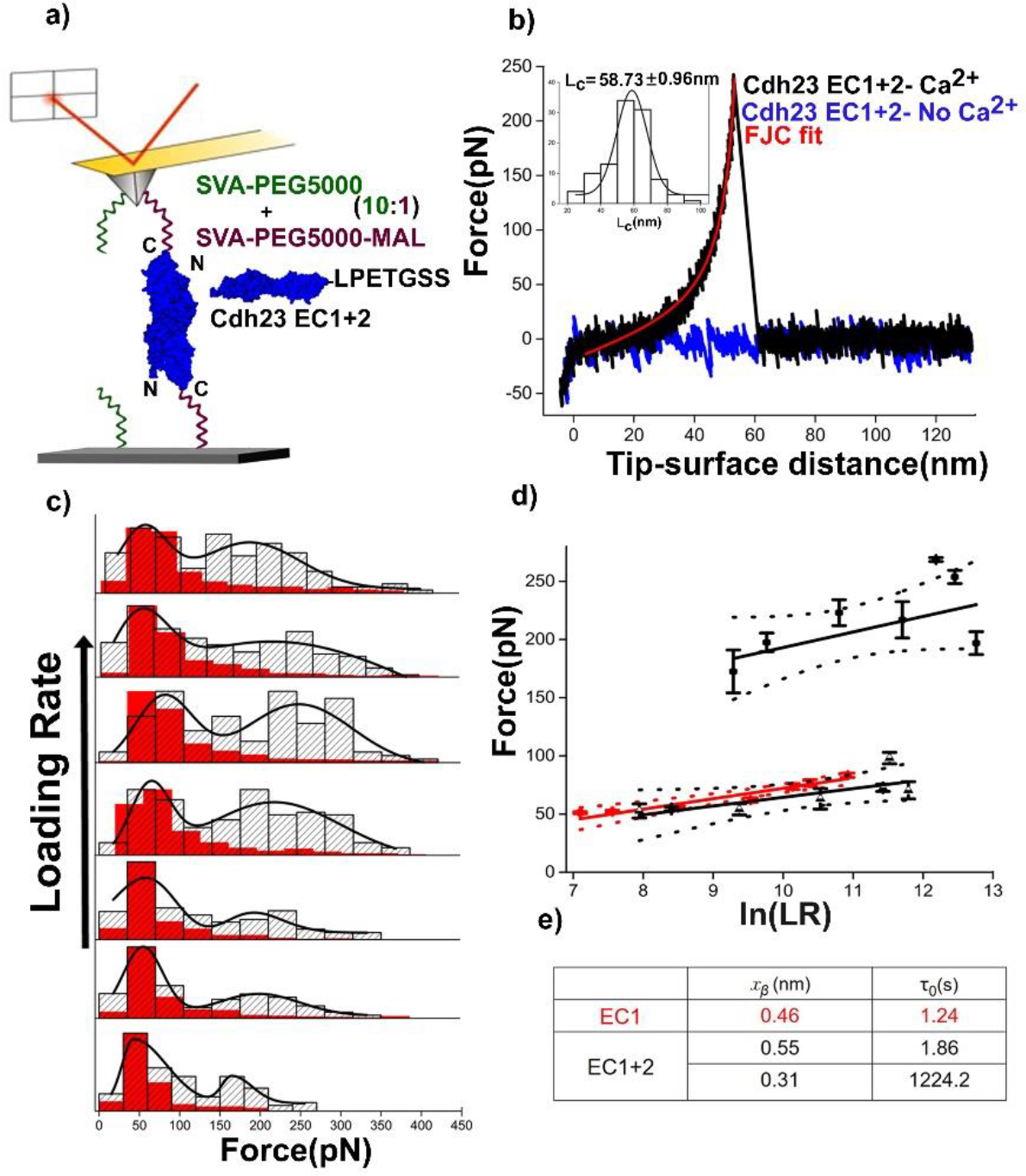
Dynamic force spectroscopy of Cdh23-EC1+2 dimers reveals two binding conformations. **a)** Schematics of the AFM cantilever and coverslip functionalized with proteins are shown for the single-molecule dynamic force spectroscopy experiments. The C-terminus of the protein is covalently attached to the cantilever and coverslip using sortase enzyme chemistry. A mixture of monofunctional and bifunctional polyethylene glycol (PEG, 5 kDa) is used as a spacer to minimize non-specific and multiple events. **b)** A typical single-molecule unbinding force curve featuring a characteristic stretching of PEG is shown in the black solid line. The fit to the freely jointed chain (FJC) model is shown in red. The rupture force is estimated from the peak maximum and contour length (*L*_*c*_) from the FJC fit. The inset shows a Gaussian distribution for *L*_*c*_ for the measurements performed at a 13841 pN/s loading-rate. The mean *L*_*c*_ was obtained from the Gaussian fit (solid line). The blue line represents no unbinding events. **c)** The histograms of unbinding forces with increasing loading-rates show a bi-Gaussian distribution for Cdh23-EC1+2 (gray) and uni-Gaussian for Cdh23-EC1 (red). Solid black curves were generated from the bi-Gaussian fitting of the distribution and used as a visual guide. From the peak maxima, we obtain the two most likely forces (F_mp_) for Cdh23-EC1+2 and one for Cdh23-EC1 at each loading-rate. **d)** The dependence of F_mp_ on ln(loading-rate) is shown for Cdh23-EC1+2 (black circles for the high-force regime and black triangles for the low-force regime) and Cdh23-EC1 (red circles). Each set of data is fitted to a Bell-Evans model (superimposed solid lines). Dotted lines indicate the 95% confidence intervals of the data. **e)** The table summarizes the parameters, x_β_ (maximum distance to unbinding in the reaction coordinate) and τ_0_ (lifetime at zero force) for EC1 and EC1+2 obtained from the Bell-Evans model fit.

A typical single-molecule unbinding force curve (black line) and a no-event (blue) curve are shown in Fig. 5b. In addition, we observed multiple unbinding events and non-specific surface-surface interactions. Overall, we observed 8–10% unbinding events, 97% of which featured single unbinding force curves that fit the freely jointed chain (FJC) model (Fig. 5b). These data corroborate well with the Poisson statistics used for single-molecule sorting. We plotted the maximum unbinding forces from each single unbinding event for six different loading-rates as histograms (Fig. 5c). The contour length (*L*_*c*_) was estimated as an FJC fitting parameter for each of the individual unbinding curves, which distributed as a single Gaussian centered at 58.7 ± 1.0 nm irrespective of the pulling rate (Fig. 5b, inset). This finding is typical for the stretching of two PEG molecules of 5 kDa molecular weight in series. The unbinding force distributions for Cdh23-EC1+2, however, showed two well-separated binomial distributions (Fig. 5c). We hypothesized that the two distributions correspond to two different binding states: the low-force distribution is due to fast binding by either of the two interacting interfaces, and the high force distribution is due to a complete handshake binding between Cdh23-EC1+2 partners. For verification, we repeated the force-spectroscopy with Cdh23-EC1 alone and measured the strength of interface one. As confirmation, the unbinding force distribution obtained for EC1 alone (red, Fig. 5c) perfectly overlaid with the low-force distribution of Cdh23-EC1+2. We also measured a higher probability of events for Cdh23-EC1 (~10%) than for EC1+2 (~6%) (Supplementary Fig. 18). This may refer to a higher binding on-rate of Cdh23-EC1 than Cdh23-EC1+2.

We then plotted the most-probable forces (F_mp_) obtained from the peak-maxima of the distributions with increasing loading-rates. The loading-rate for each velocity was estimated by considering the molecular tether and cantilever in series (Materials and Methods)(Marshall *et al*, 2006),(Ray *et al*, 2007). The data were fitted to the Bell-Evans model (<u>Eq. 6</u>)(Bell, 1978),(Evans & Ritchie, 1997) and the intrinsic lifetime(τ_0_) and the minimum distance from the bound-state to the unbound state (x_β_) were determined (Fig. 5, d and e and Supplementary Fig. 19). For wild-type EC1+2, we used two F_mp_ values for two force-distributions at each loading-rate. Our results indicate a very high τ_0_ of 1224.2 s for WT EC1+2 for the high force distribution and x_β_ of 0.31 nm. The τ_0_ obtained for wild-type EC1+2 corresponds to the strongest interactions known for any cadherin proteins. The τ_0_ and x_β_ obtained for the low-force distributions are 1.86 s and 0.55 nm, respectively, which corresponded to the values obtained for Cdh23-EC1 alone with a 95% confidence interval. Next, we repeated the force-spectroscopy experiments with mutant (E78K) and measured events similar to non-specific binding (Supplementary Fig. 18) confirming that E78K mutant does not interact with each other.

## Discussion

Even though the expression of Cdh23, both in mRNA and protein level, was shown high in a large number of normal tissues and cancer tissues where Cdh23 was found localized at the cellcell junctions, not much was explored on its involvement in cell-cell junction by forming homophilic interactions. Very recently, the homophilic *trans*-binding of Cdh23 has been reported at cell-cell junctions of breast carcinoma MCF-7 cells(Apostolopoulou & Ligon, 2012), between MCF-7 and normal breast fibroblasts, neurosensory epithelial cells, and L929 cells expressing recombinant Cdh23(Siemens *et al*, 2004). Here we are reporting the molecular details of Cdh23-EC1+2 dimers in solution for the first time. Moreover, we found that the molecular profile of the dimer is different from that of classical cadherins (Supplementary Fig. 20). All known homodimers of cadherins are formed by the overlap of EC1 alone. For Cdh23-homodimer, two outermost domains are intimately involved in dimerization and form the strongest bond as measured by SMFS. This adhesion is stronger than desmosomal cadherin-mediated Ca^2+^-independent hyper-adhesion and yet Ca^2+^ dependent(Garrod *et al*, 2005). Such strong Cdh23-adhesion is relevant in the lateral-links of stereocilia during the development of their pyramidal structure on hair-cells(Cadherin 23 is a component of the transient lateral links in the developing hair bundles of cochlear sensory cells, 2005). However, the point should be noted that the strength of the dimer might vary for full-length cadherins that mediate cell-cell adhesion. Upregulation of Cdh23 in breast carcinoma MCF-7 cells and their propensity towards heterotypic adhesion with fibroblasts in early metastasis also supports the requirement of the strong Cdh23-adhesion. A similar phenomenon is known as cadherin-switching for classical cadherins where E-cadherin, during epithelial-to-mesenchymal transition (EMT) is switched to N-cadherin, a molecule with higher binding-affinity than E-cadherin(Christofori, 2003). In summary, we have revealed a novel and unique molecular mechanism of homophilic *trans*-binding of Cdh23 that may serve as a template to understand the mechanism of homophilic adhesion by long-chain non-classical cadherins, which do not follow the typical cadherin mechanism.

## Materials and Methods

### TCGA Analysis

Relative mRNA Expression (RNA Version 2 log scale) and copy number variation (CNV) of CDH23 was obtained from cBio Cancer Genomics Portal (http://cbioportal.org) The data compared includes Breast (TCGA pub) Breast (METABRIC), Breast (TCGA 2015) (Breast TCGA (NCI) and Breast (BCCRC Xenograft)

### qRT-PCR and Western Blot

Total RNA was extracted with PureZOL RNA isolation reagent (Biorad). All samples were treated with DNase I, Amplification Grade (AMPD1-1KT, SIGMA) before cDNA synthesis with iScript cDNA Synthesis Kit (Biorad). The relative expression of 5 isoforms of Cadherin-23 mRNA was measured by using the real-time PCR (CFX Real-Time PCR Detection Systems, Biorad). (Supplementary information)

### Cloning, expression, and purification

Mouse Cdh23 (NP_075859.2) extracellular (EC) repeats EC1 (Q24 to D124) and EC1+2 (Q24 to D228) was PCR amplified using primer pair FP_Main and RP_Sortag (Table 1) and cloned into NdeI and XhoI sites of the pET21a vector (Novagen, Merck). We considered the Q24 as Q1 in our experiments. These Cdh23-EC1 and EC1+2 repeats were expressed in BL21CodonPlus (DE3)-RIPL cells (Stratagene) cultured in Luria Bertani broth (Hi-Media) and grown at 37 °C to an OD_600_ of 0.6. Induction was carried out differently for EC1 and EC1+2. EC1 was given a heat shock at 42 °C for 20 min and then the temperature of the culture was brought down to 16 °C for 30 min. It was then induced with 0.2 mM IPTG and incubated at the same temperature for 18 h. EC1+2 was induced with 0.2 mM of IPTG (Hi-Media) at 30 °C for 6 h. Unlike EC1+2, EC1 was obtained in the soluble fraction.

Cells were lysed by sonication in resuspension buffer (25 mM HEPES, 50 mM KCl, 100 mM NaCl, 2 mM CaCl_2_ pH 7.5, all from Hi-Media) and centrifuged at 4 °C for 60 min at 10,000 g. For EC1, the soluble fraction was passed twice through Ni-NTA agarose (Qiagen) equilibrated with resuspension buffer. The protein was eluted with the same buffer supplemented with 100-500 mM imidazole (1 ml each). Fractions were pooled and concentrated using Amicon Ultra (Merck Millipore). For EC1+2, after sonication and centrifugation, the pellet was isolated and washed with resuspension buffer. It was then dissolved in the resuspension buffer supplemented with 8 M urea (Promega) for 2 h on rotary-spin. The suspension was then centrifuged at 10,000 g for 30 min and the supernatant was loaded onto Ni-NTA agarose (Qiagen). The protein was eluted with 1 ml each of denaturing buffer supplemented with 25 mM, 50 mM, 100 mM, and 200 mM imidazole (Hi-Media). The eluents were pooled and refolded by step dialysis against resuspension buffer with a stepwise decrement of urea in the order 6 M, 4 M, 2 M, 1.5 M, 0.5 M and 0 M, using dialysis membranes of MWCO 2,000 (Sigma Aldrich)(Sotomayor *et al*, 2012).

All purified proteins were concentrated to 1 ml and further purified using SEC on a Superdex-75 column (GE Healthcare) in 25 mM HEPES (pH 7.5), 25 mM KCl, 100 mM NaCl, 2 mM CaCl_2_). The peak fractions were collected separately and subsequently analyzed on 12% SDS-PAGE. All mutants for EC1 and EC1+2 were processed in the same way as wildtype proteins.

**Table 1:**
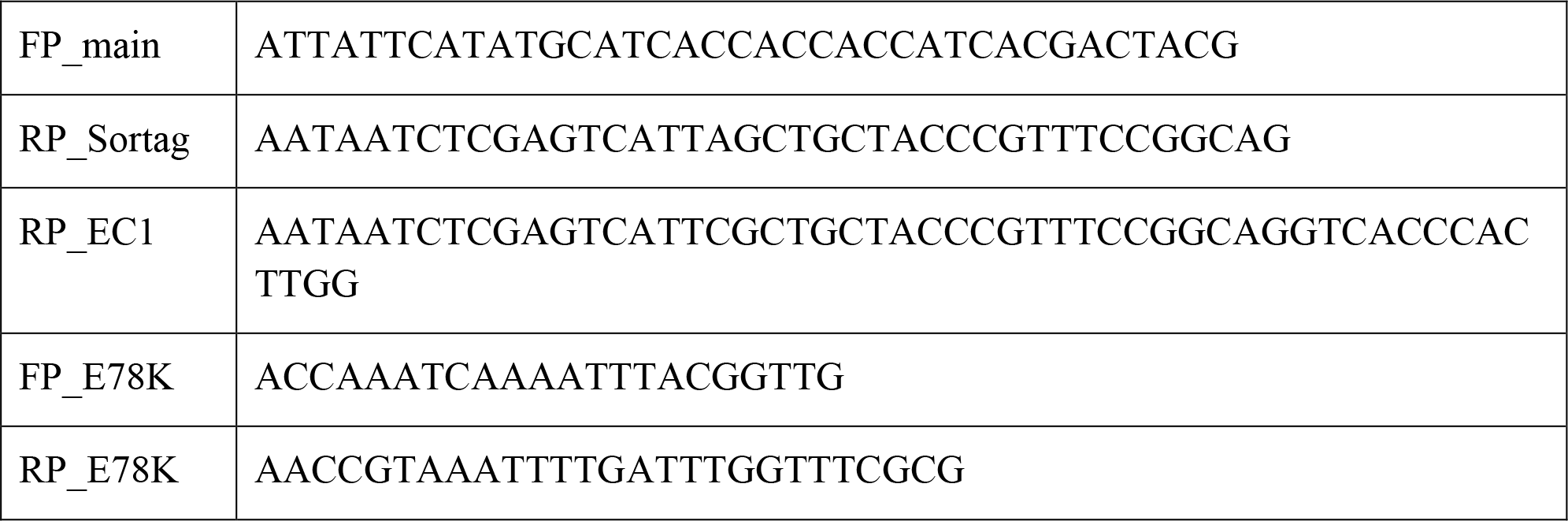
Primers used for cloning EC1, EC1+2 and mutant E78K

### Analytical ultracentrifugation

Analytical ultracentrifugation (AUC) experiments were carried out using a Beckman XLA/I ultracentrifuge, equipped with a Ti50An rotor using 12 mm six-channel cell centerpieces with sapphire windows and detection by UV at 280 nm. Both, sedimentation velocity and equilibrium experiments were performed at 20°C at pH 7.6 buffer containing 25mM HEPES, 50mM NaCl, 25mM KCl, 2mM CaCl_2_. Prior to each experiment, the protein sample was dialyzed in the buffer for 18 h.

sedimentation velocity experiments were performed at three different protein concentrations of 15 μM, 40 μM, and 56 μM, at a rotor speed of 42,000 rpm. 300 scans were taken consecutively. Data were analyzed using SEDFIT software following continuous C(s) distribution, C(M) distribution and C(s), (*f*/*f*_*0*_) models.

For equilibrium, samples were subjected to fast spins at 35,000 rpm for 20 h to achieve rapid equilibrium. Then we took one scan after each one-hour interval for five hours and checked the rmsd fluctuations to test whether samples had reached equilibrium or not. Then we decreased the rotor speed to 32,000 rpm and wait for 4 h before taking 3 scans consecutively. The Same procedure has been followed for two other speeds 28,000 rpm and 24,000 rpm respectively. For wild-type and mutant EC1-EC2 constructs, we have used two different protein concentrations, 40 μM and 60 μM respectively for equilibrium measurement. Buffer viscosity and density were measured using SEDNTERP (http://sednterp.unh.edu/). SEDPHAT was used to estimate the dissociation constant. We performed global fitting with mass conservation following monomer-dimer association model keeping baseline, meniscus, bottom, binding affinity as floating parameters.

### FRET measurements

Fluorescent dyes, cy3 as donor and cy5 as acceptor (or IAEDANS as a donor and Fluorescein maleimide as acceptor), were labeled at both the termini of protein. For N-terminal labeling, two batches of Cdh23-EC1+2 V3C of 55 μM concentration was incubated with 1:1 molar ratio of cy3 and cy5 (supplied by Lumiprobe) individually at RT for overnight. The unreacted dye was removed using spin columns (10 kDa MWCO). The labeling ratio was measured from the absorbance at 280 nm for protein, 545 nm for cy3 and 645 nm for cy5. The schematic representation for this labeling is shown in Figure1. For C-terminal labeling, cy3 and cy5 dyes were reacted with short peptide, GGGGC, at RT for overnight maintaining the labeling ratio to 1:1. Thereafter, each mixture was separately reacted to Cdh23-EC1+2 (55 μM) along with Sortase A enzyme in 4:5 ratio. With the help of sortagging chemistry, the fluorescent dye was attached to the protein. The labeled Cdh23-EC1+2 was purified from the reaction mixture using Ni-NTA where Sortase A with His-tag was bound to resin and Cdh23-EC1+2 was collected from flow-through. Labeled protein was further concentrated to 55 μM and labeling ratio was measured as 1:1. The scheme for this labeling is shown in Fig. 1d.

The absorbance spectra and emission spectra for all the four batches of proteins were measured using Cary 60 UV-Vis (Agilent technologies) and JobinYvon Fluoromax-4 Spectrofluorimeter, respectively. The possible conformations for dimers are shown pictorially in Fig. 1e: Sample 1 corresponds to NC conformation, where cy3-Cdh23-EC1+2 (N-terminus modified) and Cdh23-EC1+2-cy5 (C-terminus modified) were mixed together. Sample 2 corresponds to NN conformation, where cy3-Cdh23-EC1+2 (N-terminus modified) and cy5-Cdh23-EC1+2 (N-terminus modified) were mixed together. Sample 3 corresponds to CC conformation where Cdh23-EC1+2-cy3 (N-terminus modified) and Cdh23-EC1+2-cy5 (C-terminus modified) were mixed together. The three mixtures were incubated overnight. Emission for all the three samples along with cy3-Cdh23-EC1+2(alone) was collected from 345 nm to 650 nm with a slit width of 2 nm by exiting at 545 nm with an integration of 0.05 s and a step size of 0.1 nm.

### Pull-down down experiments at single molecule

Pull down of proteins at the single molecule level was performed on TIRF microscope. At first, the coverslip was covalently modified with unlabeled protein, Cdh23-EC1+2 WT or mutants, and then was incubated with same proteins labeled with cy3-dye at C-terminus using sortagging for 30 min (Supplementary Fig.18). Each sample was washed thoroughly to remove the nonspecific attachment. The surface was then checked for a positive signal. Images were processed by keeping the z-intensity constant for all the samples and also attaining a pseudo color for the fluorescent spots. Then, the Ca^2+^ was stripped off from the surface by washing with 1 mM EGTA (Supplementary Fig.19) and again the fluorescence was monitored. Imaging was performed using IX83 P2ZF inverted microscope (Olympus) combined with IX3 TIR MITICO TIRF illuminator equipped with 532-nm diode laser system for cy3 excitation. Fluorescence was collected using an oil-immersion objective (60X, NA 1.45, Olympus) is split into two channels by a dichroic beam splitter and recorded by electron-multiplying charge-coupled device (EMCCD) camera (Q-Imaging Roller Thunder). The filters used are Quad-band LF405/488/532/635-A-000 Bright Line Full Multi-Band Laser Filter set. Image acquisition and processing were performed using CellSens Dimension (Olympus) software.

### Live-Cell binding assay

To check whether the homophilic interactions of Cdh23 is physiologically relevant, cell-binding experiments were performed with proteins coated on coverslips. Different coverslips were covalently coated with wildtype and mutant (E78K) on the surfaces using the sortagging chemistry. After thorough washing, these coverslips were incubated with ~ 1 million MCF-7 cells for 45 min. The coverslips were then gently washed twice for 5 min with HEPES-Ca^2+^ buffer and imaged for assessing cell density. The Ca^2+^ from the surface was chelated by incubating with 1 mM EGTA and again the cell density was monitored.

### SAXS data acquisition and analysis

The SAXS data was acquired for a q-range of 0.1 to 4 nm^−1^ SAXSpace instrument (Anton Paar GmbH, Austria). The X-ray scattering setup had a slit collimated X-ray source with a wavelength of 0.154 nm. The data was collected on a Mythen (Dectris) detector placed at a distance of 317.6 mm from the sample for 60 min (20 min X 3 frames). SAXStreat software was used to calibrate the data for the beam position. The SAXSquant software was then used to subtract buffer contribution, set the usable q-range, and desmear the data using the beam profile. For each experiment, 100 μl of protein solution (Cdh23-EC1+2 and its mutant E78K) and their corresponding buffers were exposed to X-rays in a quartz capillary at a temperature of 10 °C. Data processing provided the scattering intensity (I) as a function of momentum transfer vector q (q = 4πsinθ/λ where θ and λ are the scattering angle and the X-ray wavelength respectively). The normalized Kratky plots (I(q)×q^2^ (Q × R_g_)^2^/I(0) vs. q×R_g_)) were made from the SAXS data using the program Scatter (http://www.bioisis.net/) to interpret whether the protein remains folded during the SAXS data collection. The Guinier approximation was carried out using PRIMUSQT(Konarev *et al*, 2003) of ATSAS 2.7 suite of programs(Petoukhov *et al*, 2012) in order to estimate the Radius of Gyration (R_G_) of the major scattering species.

Using GNOM program(Svergun, 1992) we carried out the indirect Fourier transformation of SAXS data to obtain the probability distribution of the pairwise vectors (P(r) curve) arising from scattering of the protein molecule in solution. The P(r) curve analysis was done to acquire the maximum linear dimension (D_max_) and the R_g_ in real space.

### Shape reconstruction

Ten independent models were generated using DAMMIF(Franke & Svergun, 2009) program.

The models were aligned, averaged and filtered using DAMAVER(Volkov & Svergun, 2003) suite of programs. The averaged envelope was further refined using DAMMIN(Svergun, 1999) program. This procedure provided an envelope that reflected the shape of a protein molecule in solution.

### Protein docking

Patch Dock is an online tool that follows rigid body docking optimization with shape-complementarity and two-point interactions between hot-spots. Hot-spots are decided based on residues that are conserved in protein-protein interaction surfaces and mediate salt-bridge type interactions, H-bonding, hydrophobic interactions, aromatic pi-stacking etc.

The available crystal structure of monomer Cdh23-EC1+2 was docked using Patch dock server(Schneidman-Duhovny *et al*, 2005) to generate the homo-dimer. Z-test was performed using SAXS based parameters like linear dimension and R_g_ and FRET-based end-end distances between the docked models and SAXS based envelop (Supplementary Table 7). The structures which agreed the most with the SAXS based envelope of the protein were overlaid by computationally aligning using SUPCOMB(Kozin & Svergun, 2001) program. Program PyMOL(Schrödinger, LLC, 2015) was used for graphical analysis and figure generation.

### Steady-state fluorescence and time-resolved anisotropy experiments

Two different concentrations of protein samples (44 μM and 11 μM) corresponding to dimer and monomer respectively were used for these experiments using JobinYvon Fluoromax-4 spectro-fluorimeter equipped with a PMT detector. In order to probe the fluorescence properties of Trp, the samples were excited at 295 nm and the emission spectra were recorded for 310-450 nm (λ_em_= 342 nm) at 25 °C. The slit-width (2 nm), step size (0.1 nm) and integration time (0.05 s) were maintained for all experiments. As a control, the Ca^2+^ ions from the dimer sample were chelated out by washing with EGTA (Sigma-Aldrich) and subsequent removal of EGTA was done by dialysis using 3,000 MWCO dialysis membrane (Sigma-Aldrich).

Fluorescence anisotropy decay measurements (for proteins of 44 μM & 11μM) were performed using time-correlated single photon counting system (Fluorocube, Horiba Jobin Yvon, NJ). For decay measurements, the emission polarizer was set to 0° with respect to excitation polarizer for parallel measurements and at 90° for perpendicular measurements. A 293 nm laser diode was used as an excitation source and the emission monochromator for tryptophan was fixed at 342 nm, at a slit width of 8 nm. The instrument response function was calculated by measuring the decay using 2% LUDOX (Sigma-Aldrich) and deconvoluting the same from the sample decay using the software provided with the instrument. A peak difference of 10,000 counts was set between the parallel and perpendicular measurements. The anisotropy decay was calculated using equation 2 and was fitted to bi-exponential decay fit to determine the values of rotational correlation times using the equation 3. The values are tabulated in SI Supplementary Table 6.

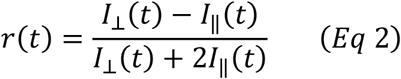

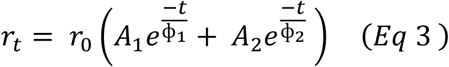

where *I*_⊥_ is the vertical emission and *I*_‖_ is the horizontal emission.

### MD Simulations

MD simulations were performed on the in-house workstation. The crystal structure of the docked model which agreed the most with the SAXS based envelop was used for simulations. Simulations were performed with GROMACS 5.0.1 using All-atom OPLS force-field and TIP4P water model. All analyses were performed using VMD. For MD simulations, the Cdh23-dimer model was placed at the center of a 13.6×5.7×5.8 nm tri-clinic box filled with four-point charge water molecules such that no atom of the protein was closer than 1 nm from the walls of the box. The MD equilibration system consisted at an average of 54,435 atoms. System charge neutrality of the system was maintained by adding Na^+^ counter-ions to the box as needed with buffer ions (50 mM NaCl, 50 mM KCl 2 mM CaCl_2_). To ensure that the solvated Cdh23-dimers model have no steric clashes or inappropriate geometries, we first energy minimized the system. Periodic boundary conditions were assumed in all simulations. A cutoff of 1 nm was used for *van der Waals* interactions. Electrostatic interactions were calculated with a particle mesh technique for Ewald summations with a 1 nm cutoff. We equilibrated the water molecules and ions around the Cdh23-dimer model in two phases. In the canonical (NVT) phase, the system was established at a constant reference temperature of 300 K. The pressure of the system was then stabilized under isothermal-isobaric (NPT) conditions. Following equilibration, 100 ns MD simulations were run at 2 fs integration steps and frames were recorded at 1 ps interval. We repeated the simulations 3 times with the same docked-structure and obtained overlapping results. We Visual Molecular Dynamics (VMD) for H-bond analysis.

### Single molecule force-spectroscopy experiments

For single molecule force-spectroscopy using AFM (Nano wizard 3, JPK Instruments, Germany), the protein molecules were immobilized on the glass coverslip and Si_3_N_4_ cantilever (Olympus, OMCL-TR400PSA-1), using a high-specificity immobilization protocol. The surfaces were cleaned with piranha (H_2_SO_4_ : H_2_O_2_ in a ratio of 3:1) (Merck) for 3 h and then incubated in 2% APTES (Sigma-Aldrich) (in 95% Acetone) and cured at 110 °C for 1 h. The amine exposed surfaces were reacted with a mixture of 10% NHS-PEG-maleimide in NHS-PEGm (LaysanBio). Both the kinds of PEG are individually dissolved in PEGylation buffer of pH 8 and then mixed to get a final concentration of 10% NHS-PEG-maleimide in NHS-PEGm. The mixture controls the density of single molecules on surfaces. A final concentration 100 μM polyglycine (GGGGC) was incubated on the surface at 25 °C for 7 h. The peptide is now attached to surface as the maleimide group on the surface reacts with cysteine of the peptide through click chemistry. This polyglycine serves as a nucleophile for sortagging chemistry. To attach the proteins on the surface, the modified surfaces are incubated with sortase A and protein (modified with LPETGGS tag at C-terminus) in 4:5 volume ratio at 200 nM concentration for each.

For quantitative estimation of force from each experiment, the spring-constant of the cantilever was measured from the thermal noise using thermal fluctuation methods(Hutter & Bechhoefer, 1993). For dynamic force spectroscopy, numerous force-distance curves were recorded at different pulling rates (500 nm/s, 750 nm/s, 2000 nm/s, 5000 nm/s, 7500 nm/s, 10000 nm/s and 15000 nm/s) while keeping approach and retract distance of 200 nm at 6 kHz sampling rate and a contact time of 500 ms constant. A total of around 6000 curves were recorded at each velocity.

The analyses for the plotted force curves were done in MATLAB with home-written programs. The single molecule events were selected from the fit to freely joint chain (FJC) model using the following equation:

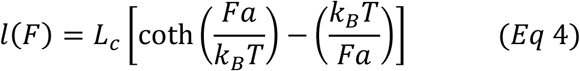

where *a* is Kuhn length, *L*_*c*_ is contour length(CL) and *l*(*F*) is stretching of PEG at every force. The unbinding forces were estimated from the peak-maxima of the single molecule unbinding events for each loading-rates and plotted as distribution. Bin-width was estimated from Scott’s method(Scott, 1979). The unbinding force-distribution at varying loading-rates showed bi-Gaussian for Cdh23-EC1+2 and single-Gaussian for Cdh23-EC1. Each force distribution was hence fitted to Gaussian distribution and the most probable force (*F*) for each loading-rate was obtained from the fit. Loading-rate at each velocity was calculated using the following equation(Ray *et al*, 2007),(Marshall *et al*, 2006)

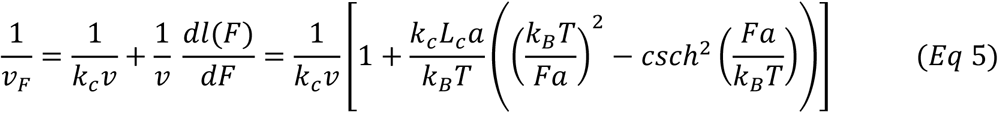

where *a* is Kuhn length, *v* is pulling velocity, *k*_*c*_ spring constant of the cantilever, *L*_*c*_ is the contour length, *v*_*F*_ loading-rate.

The most probable force (*F*) with loading-rate (*v*_*F*_) was fitted to Bell-Evans model(Bell, 1978),(Evans & Ritchie, 1997) and estimated the kinetic parameters like off-rate (*k*_0*ff*_), transition distance (*x*_β_) using the equation :

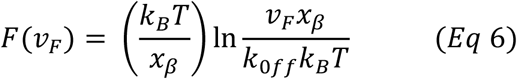

Non-specific binding rates were estimated in two different ways: by modifying either of the two surfaces (the cantilever or the coverslip) identically but without attaching proteins or by performing the same single molecule protein-protein unbinding experiments in absence of Ca^2+^ ions in the chelex buffer and EGTA buffer. In both the cases, the non-specific events accounted for <0.5% of the total number of total selected PEG stretching events.

## Supplementary Materials

Supplementary materials are attached separately.

## Acknowledgements

This work was supported by the Wellcome Trust/DBT India Alliance Fellowship [grant number: IA/I/15/1/501817] awarded to SR.

We thank Professor James W. Nelson, Stanford University, and Professor William Weis from Stanford University for critically commenting on this manuscript which certainly enhanced the quality of the work.

S.R. acknowledges the financial support by the Wellcome Trust/DBT Intermediate fellowship by India Alliance and Indian Institute of Science Education and Research Mohali (IISERM). G.S.S, A.S. and J.P.H sincerely thank IISERM for financial support. A.K. is grateful to DST INSPIRE for funding. J.S.S. is thankful to the Centre of Excellence (COE) in Frontier Areas of Science and Technology (FAST) program of the Ministry of Human Resource Development, Government of India for financial support. M.K.S. thanks to the Wellcome Trust/DBT India Alliance.

### Author Contribution

S.R. has supervised the project. G.S.S. and J.S.S. did the cloning, expression, and purification. G.S.S., A.K., A.S., A., and S.R. recorded and analyzed SAXS. J.P.H. recorded and analyzed AUC. G.S.S., A.K., A.S., R.M.Y, and S.R. analyzed MD. G.S.S. performed FRET, cell-aggregation, CD, SDS-PAGE, WB. M.K.S. performed all the qRT-PCR and TCGA analysis. G.S.S., A.S., M.K.S., and J.P.H. made the figures. S.R. wrote the manuscript. G.S.S., A.K., J.S.S., J.P.H., A.S., A., M.K.S. R.M.Y., and S.R. edited the manuscript.

A.K. and J.S.S. are co-second authors and A.S. and J.P.H. are co-third authors.

The Authors declare no competing financial interests.

## References

Apostolopoulou M & Ligon L (2012) Cadherin-23 mediates heterotypic cell-cell adhesion between breast cancer epithelial cells and fibroblasts. PLoS One 7: 1–9

Bajpai S, Correia J, Feng Y, Figueiredo J, Sun SX, Longmore GD, Suriano G & Wirtz D (2008) alpha-Catenin mediates initial E-cadherin-dependent cell-cell recognition and subsequent bond strengthening. Proc. Natl. Acad. Sci. U. S. A. 105: 18331–18336

Bell GI (1978) Models for the specific adhesion of cells to cells. Science 200: 618–27

Bork JM, Peters LM, Riazuddin S, Bernstein SL, Ahmed ZM, Ness SL, Polomeno R, Ramesh A, Schloss M, Srisailpathy CR, Wayne S, Bellman S, Desmukh D, Ahmed Z, Khan SN, Kaloustian VM, Li XC, Lalwani A, Bitner-Glindzicz M, Nance WE, et al (2001) Usher syndrome 1D and nonsyndromic autosomal recessive deafness DFNB12 are caused by allelic mutations of the novel cadherin-like gene CDH23. Am. J. Hum. Genet. 68: 26–37

Brasch J, Harrison OJ, Honig B & Shapiro L (2012) Thinking outside the cell: How cadherins drive adhesion. Trends Cell Biol. 22: 299–310

Cadherin 23 is a component of the transient lateral links in the developing hair bundles of cochlear sensory cells (2005) Dev. Biol. 280: 281–294

Christofori G (2003) Changing neighbours, changing behaviour: cell adhesion molecule-mediated signalling during tumour progression. EMBO J. 22: 2318 LP–2323

Ciatto C, Bahna F, Zampieri N, VanSteenhouse HC, Katsamba PS, Ahlsen G, Harrison OJ, Brasch J, Jin X, Posy S, Vendome J, Ranscht B, Jessell TM, Honig B & Shapiro L (2010) T-cadherin structures reveal a novel adhesive binding mechanism. Nat. Struct. Mol. Biol. 17: 339–347

Evans E & Ritchie K (1997) Dynamic strength of molecular adhesion bonds. Biophys. J. 72: 1541–55

Franke D & Svergun DI (2009) DAMMIF, a program for rapid ab-initio shape determination in small-angle scattering. J. Appl. Crystallogr. 42: 342–346

Garrod DR, Berika MY, Bardsley WF, Holmes D & Tabernero L (2005) Hyper-adhesion in desmosomes: its regulation in wound healing and possible relationship to cadherin crystal structure. J. Cell Sci. 118: 5743–5754

Gorski M, Tin A, M G, GM M, AY C, BO T, C P, A T, DI C, J C, P H, J T, M W, T A, G E, V G, TB H, LJ L, AV S, BD M, et al (2015) Genome-wide association study of kidney function decline in individuals of European descent. Kidney Int. 87: 1017–1029

Guimaraes CP, Witte MD, Theile CS, Bozkurt G, Kundrat L, Blom AEM & Ploegh HL (2013) Site-specific C-terminal and internal loop labeling of proteins using sortase-mediated reactions. Nat. Protoc. 8: 1787–99

Harrison OJ, Bahna F, Katsamba PS, Jin X, Brasch J, Vendome J, Ahlsen G, Carroll KJ,Price SR, Honig B & Shapiro L (2010) Two-step adhesive binding by classical cadherins. Nat. Struct. Mol. Biol. 17: 348–57

Hutter JL & Bechhoefer J (1993) Calibration of atomic-force microscope tips. Rev. Sci. Instrum. 64: 1868–1873

Kazmierczak P, Sakaguchi H, Tokita J, Wilson-Kubalek EM, Milligan R a, Müller U & Kachar B (2007) Cadherin 23 and protocadherin 15 interact to form tip-link filaments in sensory hair cells. Nature 449: 87–91

Konarev P V, Volkov V V, Sokolova A V, Koch MHJ & Svergun DI (2003) {\it PRIMUS}: a Windows PC-based system for small-angle scattering data analysis. J. Appl. Crystallogr. 36: 1277–1282

Kozin MB & Svergun DI (2001) Automated matching of high- and low-resolution structural models. J. Appl. Crystallogr. 34: 33–41

Lagziel A, Ahmed ZM, Schultz JM, Morell RJ, Belyantseva IA & Friedman TB (2005) Spatiotemporal pattern and isoforms of cadherin 23 in wild type and waltzer mice during inner ear hair cell development. Dev. Biol. 280: 295–306

Manibog K, Sankar K, Kim S-A, Zhang Y, Jernigan RL & Sivasankar S (2016) Molecular determinants of cadherin ideal bond formation: Conformation-dependent unbinding on a multidimensional landscape. Proc. Natl. Acad. Sci. 113: E5711–E5720

Marshall L, Helgadóttir H, Mölle M & Born J (2006) Boosting slow oscillations during sleep potentiates memory. Nature 444: 610–613

Michel V, Goodyear RJ, Weil D, Marcotti W, Perfettini I, Wolfrum U, Kros CJ, Richardson GP & Petit C (2005) Cadherin 23 is a component of the transient lateral links in the developing hair bundles of cochlear sensory cells. Dev. Biol. 280: 281–294

Panorchan P (2006) Single-molecule analysis of cadherin-mediated cell-cell adhesion. J. Cell Sci. 119: 66–74

Petoukhov M V., Franke D, Shkumatov A V., Tria G, Kikhney AG, Gajda M, Gorba C, Mertens HDT, Konarev P V. & Svergun DI (2012) New developments in the ATSAS program package for small-angle scattering data analysis. J. Appl. Crystallogr. 45: 342–350

Rakshit S, Zhang Y, Manibog K, Shafraz O & Sivasankar S (2012) Ideal, catch, and slip bonds in cadherin adhesion. Proc. Natl. Acad. Sci. 109: 18815–18820

Ray C, Brown JR & Akhremitchev BB (2007) Correction of systematic errors in singlemolecule force spectroscopy with polymeric tethers by atomic force microscopy. J.Phys. Chem. B 111: 1963–1974

Schneidman-Duhovny D, Inbar Y, Nussinov R & Wolfson HJ (2005) PatchDock and SymmDock: Servers for rigid and symmetric docking. Nucleic Acids Res. 33: 363–367

Schrödinger, LLC (2015) The {PyMOL} Molecular Graphics System, Version~1.8

Schuck P (2003) On the analysis of protein self-association by sedimentation velocity analytical ultracentrifugation. Anal. Biochem. 320: 104–124

Scott DW (1979) On optimal and data-based histograms. Biometrika 66: 605–610

Siemens J, Lillo C, Dumont R a, Reynolds A, Williams DS, Gillespie PG & Müller U (2004) Cadherin 23 is a component of the tip link in hair-cell stereocilia. Nature 428: 950–955

Sivasankar S, Zhang Y, Nelson WJ & Chu S (2009) Characterizing the Initial Encounter Complex in Cadherin Adhesion. Structure 17: 1075–1081

Söllner C, Rauch G-J, Siemens J, Geisler R, Schuster SC, Müller U & Nicolson T (2004) Mutations in cadherin 23 affect tip links in zebrafish sensory hair cells. Nature 428: 955–959

Sotomayor M, Weihofen W a, Gaudet R & Corey DP (2012) Structure of a force-conveying cadherin bond essential for inner-ear mechanotransduction. Nature 492: 128–32

Sotomayor M, Weihofen WA, Gaudet R & Corey DP (2010) Structural Determinants of Cadherin-23 Function in Hearing and Deafness. Neuron 66: 85–100

Srinivasan S, Hazra JP, Singaraju GS, Deb D & Rakshit S (2017) ESCORTing proteins directly from whole cell-lysate for single-molecule studies. Anal. Biochem. 535: 35–42

Svergun DI (1992) Determination of the regularization parameter in indirect-transform methods using perceptual criteria. J. Appl. Crystallogr. 25: 495–503

Svergun DI (1999) Restoring low resolution structure of biological macromolecules from solution scattering using simulated annealing. Biophys. J. 76: 2879–2886

Uhlén M, Fagerberg L, Hallström BM, Lindskog C, Oksvold P, Mardinoglu A, Sivertsson Å, Kampf C, Sjöstedt E, Asplund A, Olsson I, Edlund K, Lundberg E, Navani S, Szigyarto CA-K, Odeberg J, Djureinovic D, Takanen JO, Hober S, Alm T, et al (2015 Proteomics. Tissue-based map of the human proteome. Science 347

Volkov V V & Svergun DI (2003) Uniqueness of {\it ab initio} shape determination in small-angle scattering. J. Appl. Crystallogr. 36: 860–864

